# A correlative workflow streamlines synaptic imaging by cryo-electron tomography

**DOI:** 10.1101/2025.03.01.640558

**Authors:** Thanh Thao Do, Anna Siegert, Florelle Domart, Fabienne Hahn, Christina Zeising, Sarah Muth, Constantin Pape, Kathrin Kusch, Thomas Dresbach, Silvio O. Rizzoli, Arsen Petrovic, Rubén Fernández-Busnadiego

**Affiliations:** Institute of Neuropathology, University Medical Center Göttingen, 37077 Göttingen, Germany; Institute of Anatomy and Cell Biology, University Medical Center Göttingen, 37075 Göttingen, Germany; Department of Neuro-and Sensory Physiology, University Medical Center Göttingen, 37073 Göttingen, Germany; Institute of Computer Science, Georg-August-Universität Göttingen, 37077 Göttingen, Germany; Cluster of Excellence “Multiscale Bioimaging: from Molecular Machines to Networks of Excitable Cells” (MBExC), University of Göttingen, 37077 Göttingen, Germany; Institute for Auditory Neuroscience and InnerEarLab, University Medical Center Göttingen, 37099 Göttingen, Germany; Functional Auditory Genomics group, Auditory Neuroscience and Optogenetics Laboratory, German Primate Center, Göttingen, Germany; Else Kröner Fresenius Center for Optogenetic Therapies, University Medical Center Göttingen, Germany; Faculty of Physics, University of Göttingen, 37077 Göttingen, Germany

## Abstract

Despite decades of intense research, the molecular organization of the synapse is not well understood. To address this issue, we sought to develop a method for systematic imaging of synapses by cryo-electron tomography (cryo-ET), a technology capable of mapping cellular architecture at molecular resolution. Thinning of cellular samples by cryo-focused ion beam milling is a prerequisite for high-quality cryo-ET imaging, but this process needs to be guided to the structures of interest. To allow robust synaptic targeting, we established a correlative cryo-light/electron microscopy approach by which synapses are fluorescently labeled in a minimally invasive manner, using a synthetic binder of the postsynaptic scaffold PSD-95 and antibodies against the presynaptic protein Synaptotagmin-1. Cryo-ET imaging at sites of colocalization consistently revealed excitatory synapses. Our method allows structural studies of synaptic protein complexes *in situ*, facilitating investigations of the molecular architecture of synapses.

## Introduction

Synaptic contacts are highly specialized intercellular junctions mediating the electro-chemical coupling responsible for signal propagation in the nervous system. In the presynaptic terminal, synaptic vesicles (SVs) liberate neurotransmitters by fusing with the plasma membrane at a complex membrane specialization termed the active zone (AZ) (Reshetniak and Rizzoli, 2021; Sheng and Hoogenraad, 2007; Rizo, 2022; Südhof, 2012). Neurotransmitters are liberated into the space separating the pre-and postsynaptic terminals, known as the synaptic cleft, which contains adhesion complexes that establish the synaptic contact and integrate various signaling pathways (Loh et al., 2016; Missler et al., 2012; Verpoort and Wit, 2024). In excitatory postsynaptic compartments, and directly opposing the AZ, a complex array of structural and regulatory components referred to as the postsynaptic density (PSD) creates a molecular platform that concentrates glutamate receptors, including α-amino-3-hydroxy-5-methyl-4-isoxazolepropionic acid receptors (AMPARs) and N-methyl-D-aspartate receptors (NMDARs). Synaptic structures are reinforced by their close association with glial cells, and in particular with the astrocytes that wrap around synapses to support their function (Liu et al., 2023). Dysregulation of the homeostatic pathways governing synaptic function is one of the key hallmarks observed in multiple neurodegenerative and neuropsychiatric diseases (Lima Caldeira et al., 2019; Uytterhoeven et al., 2024).

Over the last decades, the molecular organization of the synapse has been intensely studied by light and electron microscopy (EM) (Dani et al., 2010; Goncalves et al., 2020). Recently, cryo-electron tomography (cryo-ET) has emerged as a powerful technique to probe the nanoscale organization of near-native cellular samples. Cryo-ET enables 3D imaging of vitrified biological material at nanometric resolution, enabling the quantitative interpretation of cellular processes and ultimately structural determination *in situ* (Keller and Fernández-Busnadiego, 2024). Several models have been explored to study synaptic biology by cryo-ET. Subcellular fractions containing functional synaptic terminals (so-called synaptosomes) have been widely used (Fernández-Busnadiego et al., 2010; Radecke et al., 2023). Primary neurons have also been grown on EM grids, and their thin regions were directly imaged by cryo-ET (Lučić et al., 2007; Petrovic et al., 2025; Schrod et al., 2018; Tao et al., 2018). Several recent studies have combined the power of fluorescence microscopy with cryo-ET to tease out the molecular specificity of excitatory and inhibitory synapses (Gogoi et al., 2022; Lapios et al., 2025; Liu et al., 2020; Tao et al., 2018). However, the thickness of these experimental models generally exceeds the mean free path of inelastic electron scattering in vitreous ice at 300 kV (∼ 300-400 nm), causing signal deterioration and a loss of high-resolution information (Rice et al., 2018). In addition, extensive screening is often needed to find regions sufficiently thin for direct imaging. Cryo-focused ion beam (cryo-FIB) milling alleviates those limitations by thinning cellular samples down to ∼150 nm, which can be imaged at high-resolution by cryo-ET (Berger et al., 2023; Noble and de Marco, 2024; Rigort et al., 2010).

Recently, several groups have made significant progress combining cryo-FIB milling and cryo-ET imaging of synapses in cultured primary neurons and brain tissues (Glynn et al., 2024; Held et al., 2024; Matsui et al., 2024). Here, we complement these advances by developing a correlative cryo-FIB-based workflow that allows efficient imaging of natively preserved synapses in cultured primary hippocampal neurons. To enable visualization of presynaptic compartments, we adapted a fluorescent labelling approach compatible with imaging at cryogenic temperatures, and we developed procedures to robustly label the postsynaptic regions of excitatory synapses. The combination of these approaches proved pivotal for the accurate targeting of synapses for cryo-ET data collection. This, in turn, allowed us to reconstruct tomograms capturing synaptic snapshots and resolving molecular details, paving the way for systematic studies of the molecular architecture of the synapse.

## Results

### A dual-color labelling strategy for synapse identification upon focused ion beam milling

We aimed to establish a workflow for efficient and high-quality cryo-ET imaging of synapses in primary neurons. To that end, we cultured rat hippocampal neurons in the presence of astrocytes on electron microscopy (EM) grids. To obtain mature and healthy neurons with dense synaptic connections, the cultures were maintained for at least 14 days *in vitro*. We used cryo-FIB milling to produce thin (∼150 nm-thick) lamellae on neuropil-rich areas of the cultures, which were transferred to a 300 kV cryo-transmission electron microscope (cryo-TEM). Low magnification overviews of these lamellae showed a rich network of axonal, dendritic and presumably astrocytic processes (Figure S1A) and occasionally neuronal cell bodies. Synaptic vesicle clusters and mitochondria could be easily discerned in these overviews due to their characteristic shapes (Figure S1B, C). However, visual identification of synapses was challenging, resulting in extended search times and only limited numbers of synaptic tomograms.

To increase targeting efficiency, we turned to cryo-correlative and electron microscopy (cryo-CLEM), whereby fluorescent labels are used to facilitate targeting for cryo-FIB-milling and subsequent cryo-ET data collection (Pierson et al., 2024). As the typical diameter of a synapse (0.5 µm – 1 µm) is similar to the cryo-ET field of view at usual magnifications, high targeting accuracy is necessary. To achieve this, we first explored live immunofluorescence approaches, which can provide high versatility and do not require genetic manipulation. However, only extracellularly-accessible epitopes can be targeted, as membrane permeabilization procedures would compromise cellular integrity. Therefore, to fluorescently label presynaptic terminals we adapted a well-established strategy using antibodies against the luminal domain of Synaptotagmin-1 (Syt1), an abundant SV transmembrane protein acting as Ca^2+^ sensor in synchronous neurotransmitter release (Rizo, 2022). Upon exocytosis, the luminal domain of Syt1 becomes exposed on the cell surface, thereby enabling immunolabeling in living neurons (Matteoli et al., 1992). Antibodies are then internalized upon endocytosis, achieving the complete labelling of the recycling SV pool by a ∼30-60 min incubation with minimal perturbation of synaptic function (Truckenbrodt et al., 2018). For each presynaptic terminal, fluorescence intensity is proportional to the amount of SV recycling, thereby not only guiding cryo-ET data acquisition, but also reflecting the activity state of the terminal imaged.

We vitrified neurons incubated for at least 30 min with Syt1 antibodies in combination with fluorescently conjugated secondary antibodies (Figure S2A), and imaged them using a fluorescence microscope integrated in the cryo-FIB instrument. Overlaying Syt1 fluorescence with ion beam-induced images showed high density of Syt1 positive puncta in neuropil regions surrounding the cell bodies (Figure S2B-C), which we targeted for cryo-FIB milling (Figure 1A). To ensure the highest targeting accuracy, we aimed to image fluorescence on the final cryo-FIB lamellae before transferring them to the cryo-transmission microscope. This is not trivial, as high-resolution cryo-ET imaging requires lamellae no thicker than ∼150 nm, which represent a minimal portion of the cellular volume and therefore may contain only a few fluorophores per synapse. Therefore, fluorescence imaging of thin lamellae requires high labeling efficiency. Importantly, we were able to detect Syt1 fluorescent puncta in final lamellae (Figure S2G), confirming the suitability of this probe for correlative targeting. Lamellae of Syt1-labelled cells were loaded into the cryo-TEM, where low magnification lamella overviews were collected (Figure 1B) and overlayed with the fluorescence images (Figure 1C). Then, tilt-series were recorded at positions coinciding with fluorescence signals (Figure 1C-F). The majority of the reconstructed tomograms contained boutons filled with SVs, often in the presence of mitochondria (e.g. Figure 1D, F). Occasionally, we also detected bona-fide synapses (e.g. Figure 1E) containing typical hallmarks such as the presence of a well-defined synaptic cleft and SVs tethered to the active zone. We also observed non-fluorescent presynaptic structures (Figure 1B, C, yellow boxes), which may correspond to silent terminals or display weak fluorescence below our detection limits. Conversely, labeled presynaptic terminals without a clear postsynaptic partner may correspond to “orphan” synapses (Krueger et al., 2003; Ziv and Garner, 2004), or to synapses where the postsynaptic terminal was removed during cryo-FIB milling. To increase the yield of clearly recognizable synapses, we decided to use another marker targeting a prominent postsynaptic feature.

**Figure 1.**
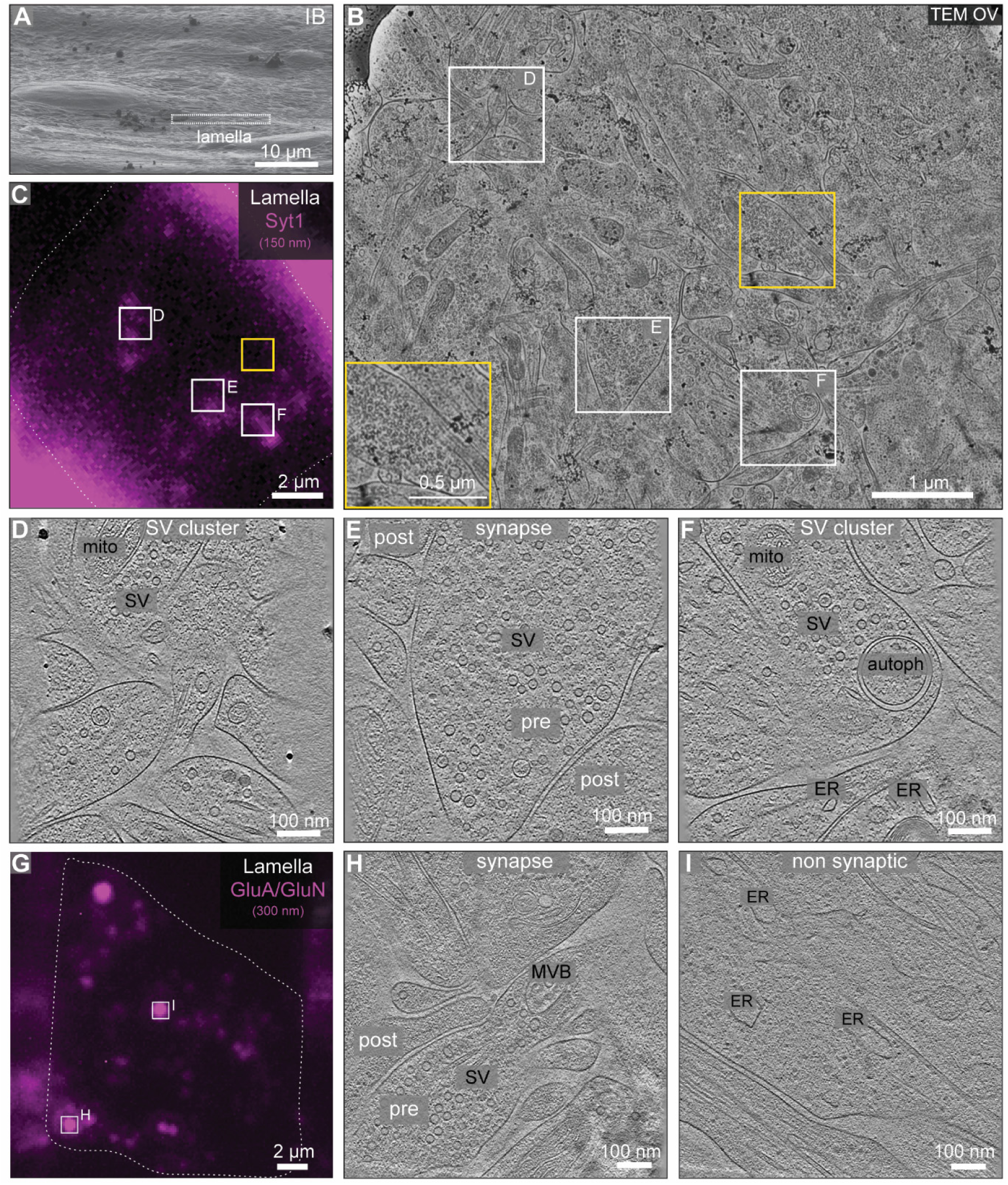
**Live immunolabelling pre-or postsynaptic terminals for correlative cryo-ET imaging. A-F**: Presynaptic targeting using presynaptic Syt1 antibodies. **A**: Ion beam-induced image of a plunge-frozen neuronal culture. A cryo-FIB lamella was produced at the boxed region. **B**: Cryo-TEM overview (OV) of the same lamella at 8700x magnification, indicating the locations of tomograms (white boxes) recorded upon correlation with the fluorescence image (C). The yellow box shows a non-labelled presynaptic terminal, magnified in the inset. **C**: Fluorescence image of the final ∼150 nm-thick lamella. Tomogram positions did not always fully match fluorescent puncta due to spatial constraints in cryo-ET imaging, such as neighboring ice crystals or the spacing required between nearby tomogram positions. **D-F**: Slices through the tomograms collected at regions indicated with white boxes in (B) and (C), displaying a synapse (E) and SV clusters (D, F). autoph: autophagosome; ER: endoplasmic reticulum; mito: mitochondrion; post: postsynaptic terminal; pre: presynaptic terminal; SV: SV cluster. **G-I**: Postsynaptic targeting using antibodies against glutamatergic receptors. **G**: Fluorescence image of a ∼300 nm-thick lamella from a neuron labelled with GluA and GluN antibodies. The lamella was milled further down to ∼150 nm thickness for tomogram acquisition at the locations marked by white boxes. **H-I**: Slices through the tomograms collected at the locations marked in (G), showing a synapse (H) and a non-synaptic region (I). MVB: multi-vesicular body.

We chose to target AMPARs and NMDARs, which reside on the excitatory PSD and are readily available for live immunolabeling (Karakas et al., 2015). Given that only 10 to 100 copies of these receptors may be present per synapse (Maynard et al., 2023; Shinohara, 2012), we reasoned that antibody labelling of both receptors with the same fluorophore would lead to an additive effect, increasing the fluorescence signal intensity in final lamellae. We used mouse GluA antibodies for AMPARs and rabbit GluN antibodies for NMDARs, together with secondary antibodies of respective species (Figure S2D) to avoid antibody-mediated cross-reactivity. As expected, we observed co-localization of both receptors with presynaptic Syt1 in room temperature immunostaining experiments (Figure S3A-B). Under cryogenic conditions before cryo-FIB milling, the combined fluorescent signal from AMPAR and NMDAR antibodies abundantly marked the neuropil regions in a similar fashion to the Syt1 antibody (Figure S2E-F). Therefore, we targeted the neuropil regions adjoining cell bodies for cryo-FIB milling. However, unlike Syt1, the GluA/GluN fluorescent signal was not preserved in the fine milled (∼150 nm thick) lamellae, and could only be reliably detected in slightly thicker (∼300-400 nm) lamellae (Figure S2H). These lamellae were later polished to ∼150 nm thickness and transferred to the cryo-TEM. Cryo-TEM overviews of the final lamellae were overlaid with the fluorescence images recorded at ∼300-400 nm lamella thickness (Figure 1G), and tilt-series were collected at fluorescent locations (Figure 1G-I). Although synaptic contacts were detected in some cases (Figure 1H), other fluorescent puncta did not correspond to synaptic regions (Figure 1I). Therefore, reliable and consistent synaptic targeting could not be achieved with postsynaptic labeling alone. To increase the targeting efficiency, an obvious strategy would be to co-label both pre-and postsynapses simultaneously. However, despite multiple attempts using secondary antibodies conjugated to different red or green fluorophores, we were not able to consistently observe signal under cryogenic conditions (Suppl Table 1).

Thus, we set out to establish an alternative strategy to target the postsynaptic compartment. We chose to label PSD-95, an abundant scaffolding component of the PSD in excitatory synapses. Although overexpression of fluorescently-tagged PSD-95 has been used to target postsynapses, this may affect important aspects of synapse physiology (El-Husseini et al., 2000; Zhang and Lisman, 2012). In contrast, fluorescent probes based on synthetic binders targeting PSD-95 allow labelling at endogenous PSD-95 levels and without any noticeable alterations in its function (Gross et al., 2013; Rimbault et al., 2024). Thus, we transduced primary hippocampal neurons at approximately DIV 5 using an adeno-associated virus carrying the recently developed PSD-95 synthetic binder Xph20-eGFP (Rimbault et al., 2024) or eGFP only as a control, and cultured the neurons until DIV 14-18. The soluble eGFP construct displayed a diffuse signal filling the entire cellular volume in room temperature immunofluorescence experiments (Figure S3C). In contrast, and in agreement with its synaptic enrichment, we observed Xph20-eGFP co-localization with the presynaptic marker Syt1 (Figure S3D). To test the applicability of Xph20-eGFP to cryo-CLEM studies, we vitrified cultured neurons simultaneously labeled with Xph20-eGFP and Syt1 (Figure S4A). Both signals were bright in cryo-CLEM, and co-localized around the cell body and along neural processes before cryo-FIB milling (Figure S4B-D). Xph20-eGFP also displayed nuclear fluorescence (Figure S4C), as unbound Xph20 is shuttled back to the nucleus to repress its own transcription upon saturation of PSD-95 binding (Rimbault et al., 2024). Most importantly, both Xph20-eGFP and Syt1 fluorescence were preserved in final ∼150 nm-thick cryo-FIB lamellae (Figure 2A). Upon transfer to the cryo-TEM, we acquired tilt-series at the positions of co-localized puncta (Figure 2A, B), which consistently revealed bona-fide synapses (Figure 2C, D). These results indicate that co-labelling of primary hippocampal neurons with Syt1 and the Xph20-eGFP PSD-95 binder leads to reliable synapse targeting for cryo-ET imaging.

**Figure 2.**
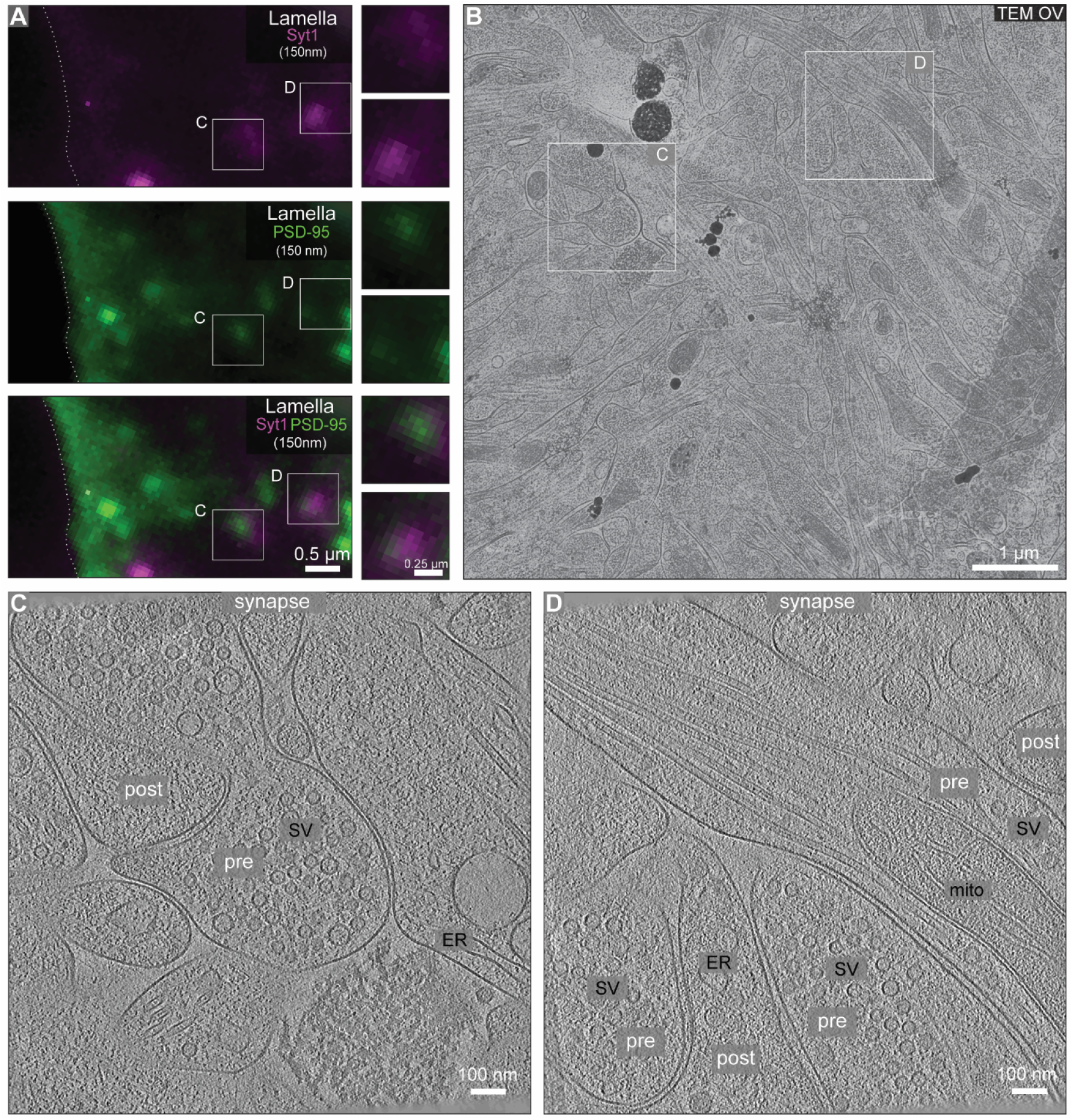
Cryo-ET on dual-color labeled of pre-and postsynaptic terminals. **A**: Fluorescence images on a 150 nm-thick lamella through a neuron co-labelled with Syt1 (magenta) and Xph20-eGFP (green). Two co-localized spots are indicated in boxes C and D. **B**: Cryo-TEM overview (OV) of the same lamella, with boxes corresponding to the spots marked in (A). **C, D**. Slices through synaptic tomograms acquired at the locations indicated in (A, B). ER: endoplasmic reticulum; mito: mitochondrion; post: postsynaptic terminal; pre: presynaptic terminal; SV: SV cluster.

### Systematic analyses of synapse morphology

The tomograms acquired on cryo-FIB milled synapses through the different labeling approaches described above showed rich ultrastructural details consistent with previous studies (Figure 3A-J). Mitochondria were often positioned in the vicinity of the SV cluster (e.g. Figure 2D and Figure 3A), which in several cases also contained microtubules traversing along (e.g. Figure 2D and Figure 3A). Within the SV cluster, individual protein densities connecting SVs (Fernández-Busnadiego et al., 2010) could clearly be discerned (e.g. Figure 3B). Molecular bridges linking SVs to the AZ membrane were also observed (Figure 3C). SV fusion intermediates were frequently visualized, consistent with the spontaneous activity reported by Syt1 labeling (e.g. Figure 3G-H). Amongst the numerous small densities present on the SV membrane, large v-ATPases could often be clearly discerned (Figure 3D), in accordance with previous *in vitro* and *in situ* studies (Coupland et al., 2024; Held et al., 2024; Kravčenko et al., 2024; Wang et al., 2024). The synaptic cleft and the PSD were populated with numerous densities of variable shape (e.g. Figure 3I). The postsynaptic compartment exhibited a diverse morphology, including cases where the cytoplasm lacked major defining structures (e.g. Figure 3A, E), and others containing various organelles such as endoplasmic reticulum, mitochondria or multivesicular bodies (e.g. Figure 3F, J).

**Figure 3:**
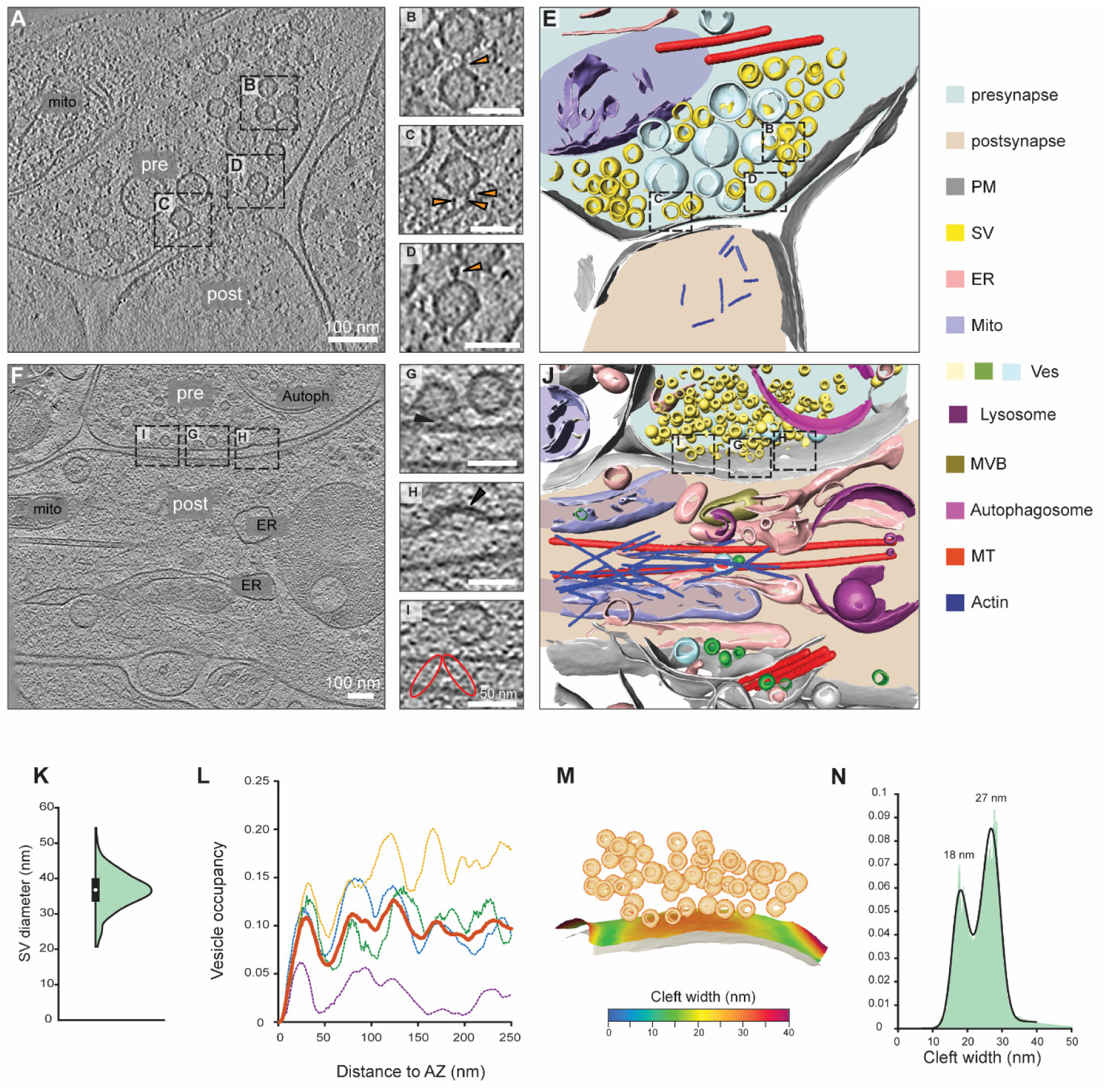
Morphological characterization of synapses upon correlative imaging. A, **F**: Slices through synaptic tomograms. autoph: autophagosome; ER: endoplasmic reticulum; mito: mitochondrion; post: postsynaptic terminal; pre: presynaptic terminal; SV: SV cluster. **E, J**: 3D segmentations of the tomograms shown in (A, F) showcasing various elements of synaptic ultrastructure (see color legend). **B-D, G-H**: magnified views with arrowheads marking the feature(s) of interest. **B**: filament connecting two SVs. **C, G**: filaments tethering SVs to the AZ. **D**: v-ATPase on the SV membrane. **H**: exo/endocytic event. **I**: synaptic cleft densities resembling pearls on a string (ellipses). **K-N**: Quantification of synaptic parameters. **K**: SV diameter (average of 37 nm ± 5.3, N=969 SVs). **L**: SV occupancy as a function of the distance to the AZ membrane. Dotted lines represent individual synapses and the bold line represents the mean. **M**: Mapping of the pre-and postsynaptic membranes distance measurements (cleft width) on the surface of the presynaptic membrane. **N**: cleft width over multiple tomograms (N=7) showing a bimodal distribution.

The average diameter of SVs (Figure 3K) in this study (37 nm) closely matched previous reports (Fernández-Busnadiego et al., 2013; Held et al., 2024). Previous studies have also shown that, under basal conditions, SVs adopt a distinct distribution within the presynaptic terminal (Fernández-Busnadiego et al., 2010; Held et al., 2024). In agreement with these data, in our tomograms the distribution of SVs displayed a characteristic peak in the proximal zone (< 45 nm from the AZ), mainly corresponding to SVs tethered to the AZ (Figure 3L). Additionally, we observed that the synaptic cleft width adopted a bimodal distribution, with a main peak at around 27 nm corresponding to the cleft central area, and a minor peak at around 18 nm associated with its periphery (Figure 3M-N). This in turn provided an objective criterion to identify synapses in cases where synapse geometry could not be visually identified in tomograms. Collectively, these measurements showcase that we can efficiently obtain cryo-ET data on natively preserved synapses, allowing subsequent molecular analyses.

### In situ structural studies of synaptic protein complexes

Large macromolecular complexes such as ribosomes were often apparent in postsynaptic compartments in our data (Figure 4A-B). Cryo-ET imaging enables structural studies of such complexes within native cellular environments using subtomogram averaging (Guo et al., 2018a; Hoffmann et al., 2022; Wagner et al., 2024; Xing et al., 2025). These procedures rely on the precise identification of a sufficient number of particles, which is often challenging due to the complex and crowded cellular environment and the imperfections of tomographic data. To test the computational detection of macromolecular complexes in our synaptic tomograms, we carried out ribosome template-matching (TM) by searching the tomograms with a low-resolution ribosome template. To visualize these results, we displayed cross-correlation values as z-score maps, whereby the cross-correlation value of each voxel was represented by the number of standard deviations separating it from the mean value of a tomogram (Cruz-León et al., 2024). TM resulted in well-defined peaks, approximately 10 standard deviations above the mean value of the tomogram (Figure 4C-D), centered at positions corresponding to manually annotated ribosomes (Figure 4B). Another prominent component of the postsynaptic compartment were clusters formed by the TRiC chaperonin (Figure 4E-F), a multiprotein complex mediating folding of major synaptic cytoskeletal proteins such as actin and tubulin and required for normal brain development (Kraft et al., 2024). We set out to investigate whether TM could detect TRiC particles by searching our synaptic tomograms with a low-resolution TRiC template, and visualizing the resulting cross-correlation as z-score maps (Figure 4G-H). TM detected individual TRiC particles consistent with manually annotated positions at ∼5 standard deviations above the mean value of the tomogram, suggesting that despite being significantly smaller than ribosomes (∼1 vs ∼2.5 MDa, respectively), TRiCs were reliably identified in postsynaptic compartments. Collectively, these analyses show that our data enables the detection of synaptic macromolecular complexes, opening the door for future systematic analyses of their *in situ* structure and (re)distribution in different synaptic states.

**Figure 4:**
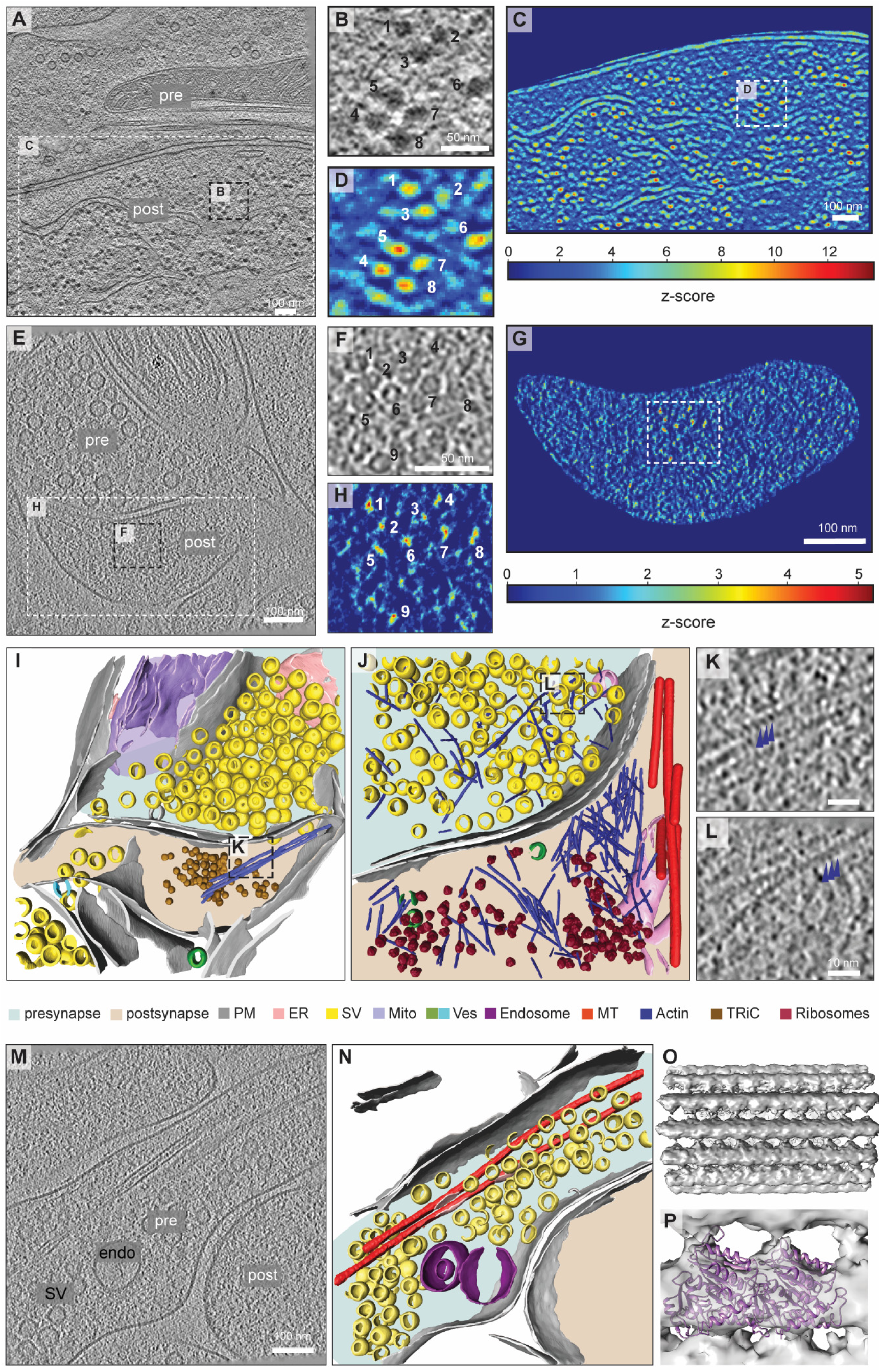
**Molecular analyses of synaptic cryo-ET data. A-G**: Template matching (TM) of ribosomes and TRiC. **A**: Slice through a synaptic tomogram with abundant postsynaptic ribosomes. The black dotted box delineates the area shown in (B, D). The white dotted box delineates the area shown in (C). **B**: magnified view of an area containing multiple ribosomes (individually numbered). **C**: Z-score heat-map of the TM results. **D**: z-score of the same region as in (B), with numbered z-score peaks corresponding to the ribosomes shown in (B). **E**: Slice of synaptic tomogram showing postsynaptic TRiC complexes. The black dotted box delineates the area shown in (F, H). The white dotted box delineates the area shown in (G). **F**: magnified view of an area containing multiple TRiCs (individually numbered). **G**: Z-score heat-map of the TM results. **H**: z-score of the same region as in (G), with numbered z-score peaks corresponding to the ribosomes shown in (G). **I, J**: 3D segmentations (see color legend) of synaptic tomograms showing synaptic actin filaments (I, J) and microtubules (J). **K, L**: magnified views of the areas shown in (I, J), showing postsynaptic (K) and presynaptic (L) actin. Arrowheads mark actin repeats. **M, N**: Slice through a synaptic tomogram (M) and its corresponding 3D segmentation (N; see color legend) showing presynaptic microtubules. SV: SV cluster, endo: endosome. **O, P**: Subtomogram averaging of presynaptic microtubules. **O**: 3D density map at ∼10.5 Å resolution. **P**: docking of a tubulin dimer (PDB 1TUB) in the density.

Several fundamental aspects of synaptic biology are critically dependent on the neuronal cytoskeleton (Gentile et al., 2022; Parato and Bartolini, 2021). While synaptic actin filaments have been extensively studied by both room temperature EM and super-resolution optical microscopy (Phillips et al., 2024), their detailed organization and dynamic rearrangements during synaptic transmission are not yet clear. In our tomograms, we visualized individual actin filaments in both pre-and postsynaptic compartments (Figure 3E, J, Figure 4I-L), with sufficient resolution to directly observe actin periodicity (Figure 4K-L). Presynaptic actin was detected less often than postsynaptic (Figure 4J, L), and typically adopted a crisscrossed organization, contrasting with the more bundled postsynaptic filaments (Figure4I-K). These data are consistent with reports on the differential regulation of pre-and postsynaptic actin pools by different cofactors (Rácz and Weinberg, 2013).

We detected microtubules in the presynaptic compartments in most of our tomograms (Figure 1H, Figure 2D, Figure 3A, Figure 4M, N), and occasionally also in postsynaptic terminals (Figure 3F, J). Previous work has studied the structure of neuronal microtubules by subtomogram averaging (Atherton et al., 2018; Chakraborty et al., 2025), but focusing on thin (< 200 nm) neuronal processes rather than synapses. Therefore, we set out to perform subtomogram averaging on presynaptic microtubules. To this end, we automatically traced presynaptic microtubules (Figure 4N, Figure S5A), oversampled the filaments and aligned the subvolumes by focusing on tubulin subunits (Figure S5B). Our final reconstruction reached ∼10.5 Å (Figure S5C), allowing confident docking of a tubulin dimer structure in our density map (Figure 4O, P). Taken together, these results confirm that the cryo-CLEM workflow presented in this study paves the way for detailed molecular analyses of synaptic components *in situ*.

## Discussion

Early synaptic cryo-ET studies focused on synaptosomes (Fernández-Busnadiego et al., 2010), a synaptic fraction prepared from brain homogenates (Whittaker, 1993). Synaptosomes are a valuable model for many aspects of synaptic function, but their isolation from brain may alter synaptic structure. Cryo-ET imaging of synapses is also possible within native tissues using vitreous sections (Fernandez-Busnadiego *et al*., 2010), or more recent cryo-FIB implementations (Glynn *et al*, 2024; Matsui *et al*, 2024). While this opens exciting possibilities, mechanistic studies require systems easily amenable to functional manipulations, such as primary neuronal cultures. Furthermore, primary cultures are also the model of choice to study the effects of mutations in key components of the synaptic machinery leading to embryonic/perinatal lethality (Kaeser *et al*, 2008; Varoqueaux *et al*, 2002; Verhage *et al*, 2000; Washbourne *et al*, 2002). Multiple cryo-ET studies have relied on cultured neurons to gain important insights into the molecular organization of excitatory and inhibitory synapses (Held *et al*, 2024; Liu *et al*, 2020; Tao *et al*, 2018). However, efficiently finding synapses suitable for high-quality cryo-ET imaging remains challenging in these samples. Some studies used correlative microscopy to guide synaptic targeting by transiently expressing fluorescently labeled synaptic proteins (Liu *et al*., 2020; Tao *et al*., 2018), but overexpression carries the risk of altering synaptic structure and function (Zhang & Lisman, 2012). Furthermore, while correlation may inform on synaptic locations, without suitable sample thinning methods most of these positions may be too thick for high-quality imaging, as synapses often form in areas of overlapping neuronal process. High-quality cryo-ET requires 100-200 nm-thick samples, which can now be efficiently prepared by cryo-FIB milling (Young and Villa, 2023). In fact, a recent study took advantage of cryo-FIB to thin down neuronal cultures and directly image synapses in the resulting lamellae (Held *et al*., 2024), but in our hands such approach led to low targeting efficiency and only small numbers of bona-fide synaptic tomograms.

Our correlative cryo-ET workflow addresses this issue. We tested an extended list of reagents aiming to fulfill two main requirements. First, the labeling should introduce minimal functional disturbances in the synapse, as cryo-ET must be performed in as close-to-native conditions as possible. At the same time, the fluorescent labels should enable precise synaptic targeting, as synapse dimensions are comparable to the cryo-ET field of view. While cryo-light microscopy imaging can also be performed in whole cells before cryo-FIB milling, this approach suffers from the limited accuracy of 3D correlation, especially in the Z direction, due to the poor resolution of the long-working distance of cryo-LM objectives (Pierson *et al*, 2024). Slight errors in lamella positioning may lead to completely missing the region of interest. Therefore, we adopted a post-milling correlative approach to maximize targeting efficiency. We used whole cell fluorescence to target synapse-rich regions for cryo-FIB milling, acquired cryo-LM on the resulting thin lamellae, and used this information to guide synaptic localization in the cryo-TEM.

The most efficient targeting approach was the simultaneous dual-color labeling with antibodies against the luminal domain of Syt1 (presynaptic) and a PSD-95 synthetic binder (postsynaptic). The Syt1 assay is well-established, and results in efficient labeling of recycling SVs with little-to-no influence on synaptic physiology (Matteoli *et al*, 1992; Truckenbrodt *et al*, 2018). Besides aiding correlative synapse targeting, this assay also informs on the recycling activity of the presynaptic terminal imaged, and may also enable future developments such as the identification of recycled SVs by immunogold labeling. Despite multiple attempts to combine Syt1 labeling with postsynaptic live immunolabeling of AMPARs and NMDARs, none of the fluorophore combinations tested was bright enough in cryogenic conditions to allow simultaneous Syt1 and GluA/GluN imaging on cryo-FIB lamellae. The reasons behind these problems require further investigation, and may involve insufficient labeling density and/or spectral shifts of the fluorophores at liquid nitrogen temperature. To circumvent these issues, we resorted to a recently developed highly specific synthetic binder of PSD-95, which was designed to minimize any possible influence of synaptic function (Rimbault *et al*, 2024) and resulted in bright fluorescence that was preserved on final cryo-FIB lamellae. The combination of Syt1 and PSD-95 fluorescent labeling enabled efficient targeting of excitatory synapses for cryo-ET imaging. Future developments will aim to extend these approaches to other synapse types, including inhibitory and dopaminergic.

Our labeling strategy is versatile, and can be combined with any functional manipulation and genetic background. In addition, this methodology can be used in conjunction with genetic code expansion and biorthogonal click chemistry approaches for live labelling of a wide range of synaptic proteins in primary neurons (Stajković et al., 2023). Thus, this workflow facilitates structure-function studies on the molecular architecture of synapses. To showcase these possibilities, we demonstrated the localization of major synaptic macromolecules using template matching, as well as subtomogram averaging of synaptic microtubules with nanometric resolution. At this moment, the major resolution-limiting factor is the size of the dataset, but the tools are now in place to systematically acquire sufficient synaptic cryo-ET data to study the still poorly-defined structural bases of fundamental phenomena like neurotransmitter release, synaptic plasticity or pathological synapse dysfunction.

## Acknowledgments

We thank Dirk Schwitters and Daniela Gerke for technical assistance, Felipe Opazo for help on immunolabeling strategies, Tat Cheng for support with cryo-ET imaging, and Tanvir Shaikh for support with data analysis. We also thank Saikat Chakraborty, Eric Hosy, Jenny Keller, Vladan Lučić and Jonathan Wagner for insightful suggestions, and all members of the Busnadiego laboratory for helpful discussions. This work was supported by the Deutsche Forschungsgemeinschaft (DFG, German Research Foundation) under Germany’s Excellence Strategy (EXC 2067/1-390729940), NeuroNex2 (FE 1940/1-1) and SFB1286 (projects A03, A12 and C10), as well as by the Else Kröner Fresenius Foundation via the Else Kröner Fresenius Center for Optogenetic Therapies. Cryo-ET instrumentation was jointly funded by the DFG Major Research Instrumentation program (448415290), the Ministry of Science and Culture of the State of Lower Saxony.

## Materials and methods

### Antibodies

The majority of the data shown in this study (Fig. 1, Fig. 2, Fig. S2, Fig. S3C-D, Fig. S4) was generated using the following primary antibodies: mouse anti-Synaptotagmin 1 luminal domain (105 311, Synaptic Systems, 1:100 dilution; hereafter “mouse anti-Syt1”), mouse anti-GluA extracellular domain (182 411, Synaptic Systems, 1:200 dilution; “mouse anti-GluA”); rabbit anti-GluN1 extracellular domain (114 103, Synaptic System, 1:200 dilution; “rabbit anti-GluN”). These secondary antibodies were used: goat anti-mouse IgG (H+L) Alexa Fluor 647 (A-21234, ThermoFisher Scientific, 1:200 dilution; “anti-mouse-AF647”), goat anti-rabbit IgG (H+L) Alexa Fluor 647 (A-21245, ThermoFisher Scientific, 1:200 dilution; “anti-rabbit-AF647”).

During the development of the method (Fig. S3A-B), we also used the following antibodies: mouse anti-Synaptotagmin1 luminal domain directly conjugated to ATTO 647N (105 311AT647N, Synaptic Systems, 1:100-1:500 dilution; “Syt1-ATTO647N”), mouse anti-GluA extracellular domain conjugated to ATTO 488 (182 411AT488, Synaptic Systems, 1:200 dilution; “GluA-ATTO488”), rabbit anti-GluN1 extracellular domain conjugated to ATTO 565 (generated by Synaptic Systems, 1: 200 dilution; “GluN-ATTO565”). In addition, the following antibodies were successfully used in room temperature staining, but did not provide sufficient signal under cryo conditions (Suppl. Table 1): rabbit anti-Synaptotagmin1 luminal domain (105 308, Synaptic Systems, 1:100 dilution, “rabbit-anti-Syt1”), goat anti-rabbit IgG (H+L) ATTO 565 (2307, Hypermol, 1:100 dilution; “anti-rabbit-ATTO565”), goat anti-mouse IgG (H+L) Alexa Fluor 488 (A11001, ThermoFisher Scientific, 1:200 dilution, “anti-mouse-AF488”).

### Preparation of primary rat hippocampal neuron cultures

All procedures followed the guideline and regulations of the University of Göttingen and the State of Lower Saxony (Niedersächsisches Landesamt für Verbraucherschutz und Lebensmittelsicherheit – LAVES, Germany).

Primary hippocampal cultures were prepared from newborn rat pups (P0-P1) following standard procedures (Banker and Cowan, 1977; Jähne et al., 2021). In brief, hippocampi were dissected in ice cold Hank’s balanced salt solution and incubated in an enzyme solution (Dulbecco’s Modified Eagle’s Medium (DMEM) supplemented with 0.2 mg/mL cysteine, 100 mM CaCl_2_, 50 mM EDTA and 20-25 U/mL papain; bubbled with carbogen for 10-20 min and preheated to 37 °C) for one hour at 37 °C on a shaker at approx. 30 rpm. Afterwards, the enzymatic reaction was stopped by incubating hippocampi in an inactivation solution (DMEM supplemented with 5 mg/mL bovine serum albumin and 10% fetal calf serum) for 15 min at 37 °C followed by three washes with washing medium (MEM supplemented with 3.3 mM glucose, 2 mM L-glutamine and 10% horse serum). Hippocampi were homogenized in washing medium, triturated with a 10 mL serological pipette, filtered through a 100 μm cell strainer then centrifuged for 8 min at 800 rpm. The supernatant was removed, leaving dissociated neurons in suspension. Neurons were seeded in plating medium (MEM supplemented with 2.5 mM glucose, 2 mM L-glutamine and 10% horse serum). After seeding on coverslips or EM grids (as described below), cells were allowed to settle for 2 h at 37°C followed by medium exchange to Neurobasal-A medium (Life Technologies, Carlsbad, CA, USA) containing 2% B27+ supplement (Gibco, ThermoFisher Scientific, USA) and 1% GlutaMax (Gibco, ThermoFisher Scientific, USA). Cultures were maintained under sterile conditions for 14-21 days at 37 °C and 5% CO_2_.

For a subset of experiments (Figure 4, Figure S1, Figure S3A), primary neurons were prepared from E19 rat embryos. The procedure resembled the preparation of postnatal cultures, with the following differences. Hippocampi were digested in 0.025% Trypsin and deactivated in Hank’s balanced salt solution. Single neurons were obtained by passing the tissue through 20G and 25G needles, then plated in DMEM supplemented with 10% fetal calf serum, 1% penicillin/streptomycin and 2 mM L-Glutamine.

### Live immunostaining

For live staining of pre-and postsynapses, the primary and secondary antibodies were pre-incubated together in growing medium (350-500 µL for grids on glass bottom dishes, 100-300 µL for coverslips) for 5 min at 37 °C. Presynaptic terminals were marked using antibodies against the luminal domain of Synaptotagmin 1 (mouse anti-Syt1, anti-mouse-AF647; Syt1-ATTO647N) for 30-60 min at 37 °C to label the entire recycling pool (Truckenbrodt et al., 2018). For the postsynaptic staining, antibodies targeting the extracellular domains of AMPA receptors (mouse anti-GluA, anti-mouse-AF647) and NMDA receptors (rabbit anti-GluN, anti-rabbit-AF647) were pre-incubated together to enhance fluorescent signal. The staining was performed either for 7 min at 37°C or for 4 min at 4°C.

### AAV production and infection

Recombinant AAV2/1 viral vectors were produced according to standard AAV purification procedures (Huet and Rankovic, 2021). In brief, triple transfection of HEK-293T cells was performed using pHelper plasmid (TaKaRa/Clontech), the trans-plasmid providing viral capsid AAV1 (Plasmidfactory, PF0401) and the cis-plasmid providing the hSyn_Xph20-eGFP-CCR5TC_WPRE_SV40pA (Addgene plasmid # 187444, a gift from Matthieu Sainlos) or hSyn_EGFP_WPRE_hGH (Addgene plasmid # 58867, a gift from Edward Boyden) transgene cassettes. Viral particles were harvested from the supernatant 72 h after transfection, and additionally from cells and the supernatant 120 h after transfection. Particles were precipitated from the supernatant using 8% [w/v] polyethylene glycol 8000 (Acros Organics, Germany) in 500 mM NaCl for 2 h at 4 °C. Both the precipitate and the cell pellet were lysed in high salt buffer (500 mM NaCl, 2 mM MgCl_2_, 40 mM Tris–HCl, pH 8.0) and non-viral DNA was degraded using salt-activated nuclease (Arcticzymes, USA). After clarification at 2000 x g for 10 min, viral vectors were purified by density step gradient centrifugation using iodixanol (Optiprep, Axis Shield, Norway) with 15, 25, 40, and 60% [v/v] at 350,000 x g for 2.25 h (Grieger et al., 2006; Zolotukhin et al., 1999). Finally, viral particles were sterile filtered and buffer was exchanged to sterile PBS supplemented with 0.001% [v/v] Poloxamer 188 (Gibco, Germany) using Amicon filters (EMD, UFC910024).

Viral vector titers were determined as the number of DNase I resistant genome copies (GC) using naica dPCR (Stilla Technologies) with naica multiplex PCR mix (Stilla Technologies) and a primer-probe set specific for ITR2 on a Ruby chip (Stilla Technologies). Primers were GTATGAGTGCAAGTGGGTTT, AACCCCTAGTGATGGAGTTG, and the probe CCTCAGTGAGCGAGCGAGC was coupled to FAM at 5’ with BHQ-1 at 3’ as quencher. Titers were 2.75E+11 GC/ml for AAV2/1_hSyn_ Xph20-eGFP-CCR5TC_WPRE_SV40pA (hereafter Xph20-eGFP) and 8.31E+10 GC/ml for AAV2/1_hSyn_EGFP_WPRE_hGH (hereafter eGFP). The purity of viruses was checked by silver staining (Pierce, Germany) after gel electrophoresis (NovexTM 4–12% Tris-Glycine, Thermo Fisher Scientific).

Primary neurons were infected with AAVs at DIV 4-8 in fresh growing medium and incubated at 37°C and 5% CO_2_ until the day of experiment (DIV 14-18). Xph20-eGFP and eGFP were transduced at a multiplicity of infection of 30,000-70,000, and 16,000, respectively.

### Immunohistochemistry

For immunofluorescence, neurons were seeded in plating medium in 24-well plates at a density of 50,000 cells per 12 mm coverslips, coated with 1 mg/mL poly-L-lysine. Cells were infected with virus or live labelled with antibodies as described above. Afterwards, cells were fixed for 20 min with 4% paraformaldehyde in PBS, followed by 10 min incubation in quenching solution (50 mM NH_4_Cl in PBS). Coverslips were mounted on glass slides using Vectashield mounting medium. Images were obtained at a DMi8 microscope (Leica) using 63x oil immersion objective and LasX software (Leica).

### Seeding on EM grids

For cryo-ET, hippocampal cells were seeded onto EM grids according to the following procedure. Four holey EM grids (R2/2, Au 200 mesh, 100 holey SiO_2_ film; Quantifoil Micro Tools GmbH) were placed into the deepening of a sterile 35 mm glass bottom dish and plasma cleaned for 30 s at medium voltage (PDC-32G-2, Harrick Plasma Cleaner). Subsequently, grids were handled under sterile conditions at all times. Grids were coated with 1 mg/mL poly-L-lysine (#P2658, Sigma Aldrich) in borate buffer (80 mM H_3_BO_3_, 20 mM Na_2_B_4_O_7_, pH 8.5) for 2 h at 37 °C followed by four washes with sterile ddH_2_O. Coated grids were stored in ddH_2_O at 4 °C until seeding (up to a week). For the seeding, ddH_2_O was removed and grids were equilibrated in plating medium for at least 1 h. Cells were pipetted onto the grids in plating medium at a density of 20,000 cells per grid.

### Plunge freezing

Immediately after immuno-live staining, cells were washed three times in Tyrode’s solution (ThermoFisher Scientific) and incubated for 2-10 min in Tyrode’s solution supplemented with 5% glycerol (v/v) to facilitate vitrification (Bäuerlein et al., 2023, 2017). For plunge freezing, grids were mounted on a lab-made manual plunger and manually blotted from the back with filter paper (Whatman, 597) for 13 s, followed by rapid plunging into a mixture of liquid ethane-propane (37% ethane, 63% propane) cooled to liquid nitrogen temperature (approx.-196 °C). Subsequently, the grids were kept in liquid nitrogen until microscope loading.

### Cryo-correlative microscopy and cryo-FIB milling

For cryo-FIB milling, samples were transferred into an Aquilos 2 cryo-FIB microscope (ThermoFisher Scientific) equipped with an iFLM integrated light microscope (ThermoFisher Scientific). Prior to cryo-FIB milling, SEM grid overviews were acquired as tile sets with approx. 4000x magnification at 3-5 kV using MAPS software 3.14 (ThermoFisher Scientific), and milling positions (i.e. neuronal cell bodies in the centre of grid squares) were localized on the SEM tile set. iFLM was used for acquisition of pre-and/or post-synaptic fluorescence at either of these stages, depending on the experiment: (1) before milling, (2) after rough milling (600 nm lamella thickness), (3) after medium milling (300 nm lamella thickness) or (4) after fine milling (150 nm lamella thickness). iFLM data was acquired as z-stacks with a range of-3 µm to +3 µm (step size 0.3 µm) for stage 1,-2 µm to +2 µm (step size 0.2 µm) for stage 2 and-1 to +1 µm (step size 0.1 µm) for stages 3 and 4. Samples were imaged in the 625 and 470 nm channels at 40-50% intensity and 500 ms. For each of the four iFLM stages, z-stacks were complemented with single images acquired in reflective light mode for correlation. Additionally, to facilitate correlation of iFLM data acquired before milling (stage 1), fiducial crosses were milled into the grid bars before iFLM acquisition. Correlation of iFLM data with ion-beam induced images of neuronal cell bodies and/or lamellae, was performed as previously described (Guo et al., 2018b) using three-point alignment in MAPS software 3.14 (ThermoFisher Scientific). To protect cells from charging damage caused by out-of-focus gallium ions during cryo-FIB milling, a layer of inorganic platinum was deposited on the grids by sputter coating (15 s, 30 mA, 0.1 mbar) followed by organic platinum deposition using a gas injection system (30 s) and another sputter coating (15 s, 30 mA, 0.1 mbar). Cryo-FIB milling was performed at 9-10° milling angle. Lamellae were milled at ion beam intensities between 1 nA (rough milling) and 30 pA (fine milling) and reached a final thickness of approximately 150 nm.

### Cryo-electron tomography

Tilt series were acquired in a 300 kV Krios G4 cryo-TEM (ThermoFisher Scientific) equipped with a Falcon 4i direct electron detector (ThermoFisher Scientific) and a Selectris energy filter (ThermoFisher Scientific). For the energy filter, a slit width of 20 eV was used during tilt series acquisition. Tilt series were recorded using PACE-tomo (Eisenstein et al., 2023) implemented within SerialEM 4.0 (Mastronarde, 2005) in a dose-symmetric fashion with tilt angles ranging from-45° to +63° with an angular increment of 3°. The total electron dose per tilt series was 120 e^−^/Å^2^. Data was collected at 33k, 42k or 64k magnification, resulting in a pixel size of 3.65 Å, 2.56 Å and 1.89 Å, respectively. Pre-processing of tilt series was automated using an in-house script (https://github.com/rubenlab/snartomo) performing the following steps: frame alignment in Motioncor2 1.6.5 (Zheng et al., 2017), CTF estimation in CTFfind 4.1.14 (Rohou and Grigorieff, 2015), and tilt series alignment and reconstruction in IMOD 4.12 (Mastronarde, 1997). Reconstruction was based on patch tracking (without fiducials) and the weighted back projection method. For visualization purposes, the respective tomograms were filtered by applying Wiener-like filter (https://github.com/dtegunov/tom_deconv).

### Tomogram segmentation

Membrane segmentation was done on bin 4 tomograms automatically in MemBrain-seg (Isensee et al., 2021; Lamm et al., 2024) and refined in Amira (Thermo Fisher Scientific). The lumen of the organelles was manually filled in. The tracing of microtubules and actin filaments was performed using the XFiber module implemented in Amira. For synaptic vesicle analysis (Figure 3K, L), synaptic vesicles were segmented using the SynapseNet software (Muth et al., 2024) using unsupervised domain adaptation.

### Synaptic vesicle analysis

SV diameter (Figure 3K) and distribution profile (Figure 3L) were calculated using Pyto (Lučić et al., 2016). SV distribution was calculated as the volume fraction of the presynaptic cytoplasm (divided into 1-voxel thick layers) occupied by SVs.

### Synaptic cleft analysis

The synaptic cleft width (Figure 3M, N) was calculated as the distance between the pre-and postsynaptic plasma membranes using the surface morphometrics pipeline (Barad et al., 2023). Results were analysed in MATLAB.

### Template matching

Template matching (TM) procedures using the ribosome (EMD-5592) and the TRiC chaperone (EMD-32922) as templates were performed using Dynamo (Castaño-Díez et al., 2012) and PyTom (Hrabe et al., 2012; Chaillet et al., 2024), respectively. The templates were low-pass filtered to 40 Å. The cross-correlation volumes were transformed to z-scores and plotted in MATLAB for visual inspection of the TM outcome (Figure 4C, D, G, H).

### Subtomogram averaging

The coordinates obtained from the XFiber tracing in Amira were used to create a tubular grid using STOPGAP (Wan et al., 2024), by placing points on each tubulin monomer (every 4 nm along the tube and every 6 nm around the tube). Subtomograms were subsequently extracted from 76 MTs from 18 bin 4 tomograms (7.56 Å/pixel). Subtomograms from one MT were aligned and averaged to create an initial reference, and all subtomograms were then aligned to this reference using STOPGAP. To determine their polarity, MTs were individually averaged using multiclass averaging. The polarity of each MT was visually inspected and flipped to create a dataset with uniform polarity. Subtomograms were then aligned and averaged first in bin 4 and then in bin2 (Figure 4P).

