## Supplementary information for "A correlative workflow streamlines synaptic imaging by cryo-electron tomography"

| Presynaptic label | Postsynaptic label | Signal detected at cryogenic temperatures before cryo-FIB milling |
|---|---|---|
| Mouse anti-Syt1<br>Anti-mouse-AF647 | Rabbit anti-GluN<br>Anti-rabbit-AF488 | Presynaptic |
| Rabbit anti-Syt1<br>Anti-rabbit-AF647 | Mouse anti-GluA<br>Anti-mouse-AF488 | Presynaptic |
| Rabbit anti-Syt1<br>Anti-rabbit-ATTO565 | Mouse anti-GluA<br>Anti-mouse-AF647 | Postsynaptic |
| Mouse anti-Syt1<br>Anti-mouse-AF647 | - | Presynaptic |
| Rabbit anti-Syt1<br>Anti-rabbit-AF647 | - | Presynaptic |
| Mouse anti-Syt1-ATTO647N | - | Presynaptic |
| Rabbit anti-Syt1<br>Anti-rabbit-ATTO565 | - | - |
| Rabbit anti-Syt1<br>Anti-rabbit-AF488 | - | - |
| - | Mouse anti-GluA<br>Anti-mouse-AF488 | - |
| - | Mouse anti-GluA<br>Anti-mouse-AF647 | Postsynaptic |

**Table S1. Detection of fluorescence of immunolabeled pre- and postsynaptic terminals at cryogenic temperatures**. All configurations displayed typical synaptic staining at room temperature (not shown), but only Alexa Fluor 647 (AF647) fluorescence was observed at cryogenic temperatures prior to cryo-FIB milling, preventing dual-colour immunolabelling.

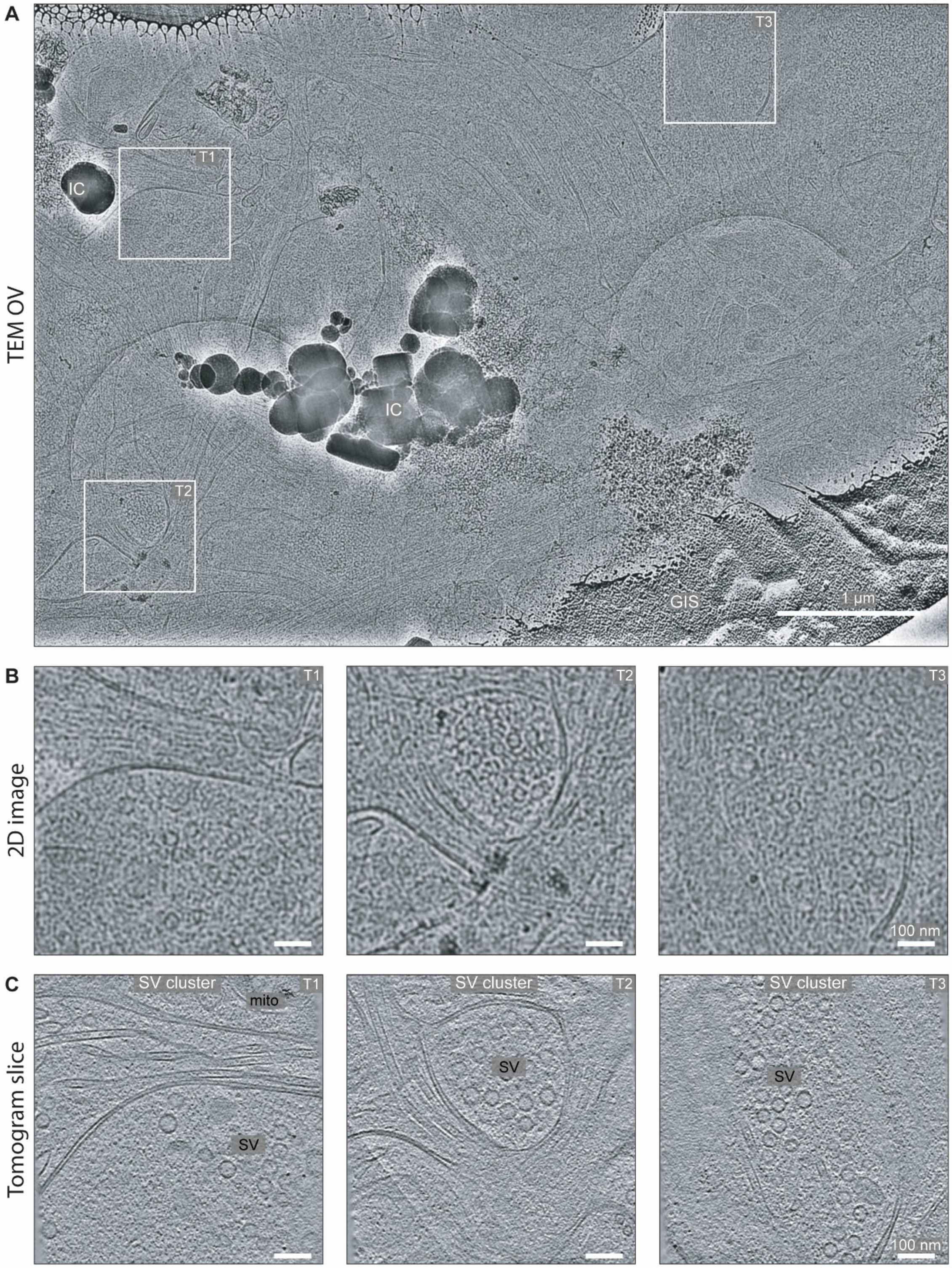


**Figure S1. Synapse targeting on non-fluorescently labeled neurons. A**: Cryo-TEM (OV) low magnification (8700x) overview, showing a complex network of neuronal processes. GIS: platinum layer deposited by the gas injection system; IC: superficial ice crystal contamination. **B**: Magnified views of regions indicated in (A) showing putative synaptic locations where tomograms were collected. **C**: Slices of tomograms recorded at regions shown in (B), containing SV clusters but no actual synapses.

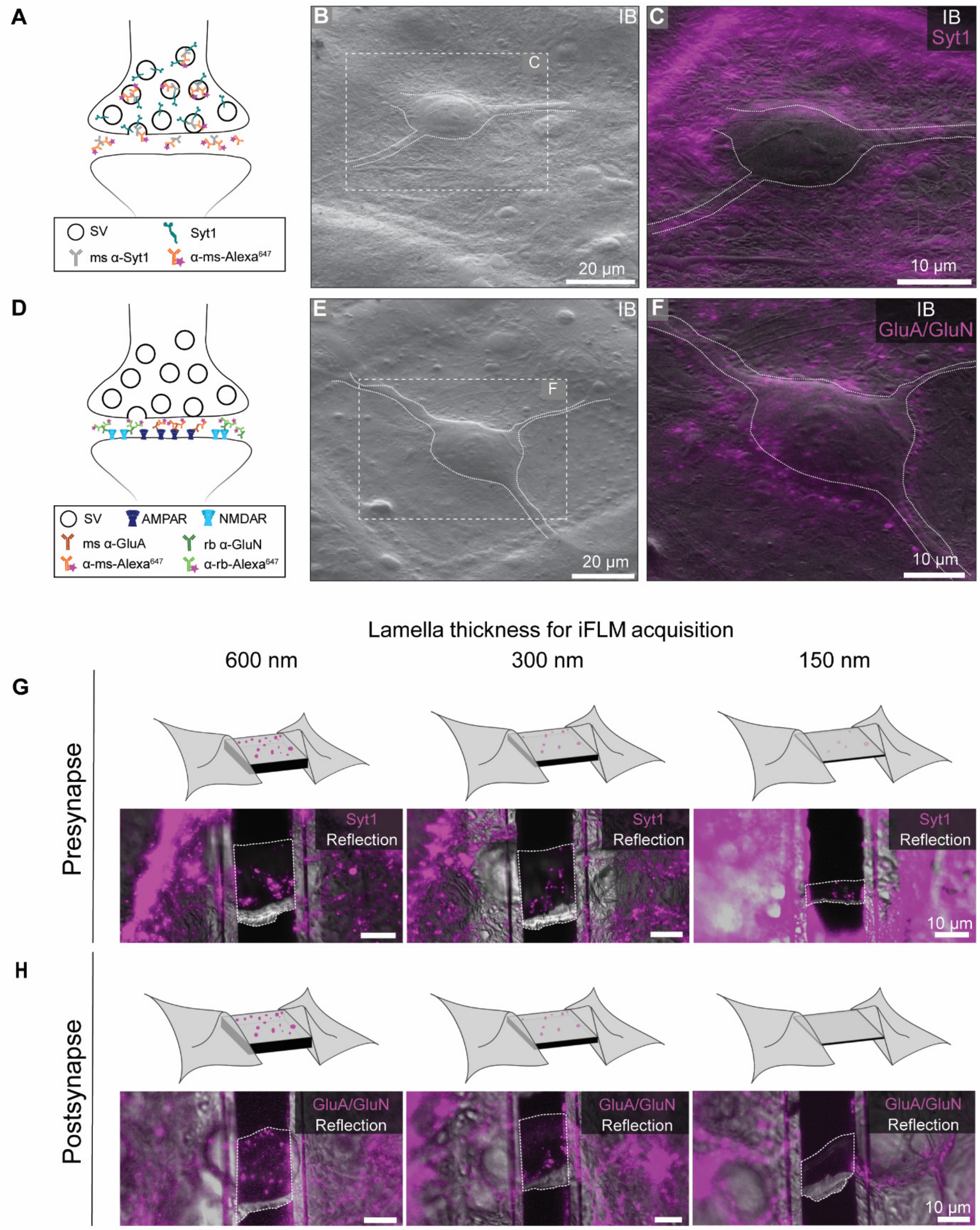


**Figure S2. Cryo-FIB milling of neurons upon live immunolabeling. A-C**: Labelling of presynaptic terminals using Syt1 antibodies (mouse anti-Syt1, anti-mouse-AF 647). **D-F**: Labelling of postsynapses using GluA and GluN antibodies (mouse anti-GluA, anti-mouse-AF647; rabbit anti-GluN, anti-rabbit-AF647). **A, D**: Cartoons depicting the antibodies used and their cellular targets. **B, E**: Ion beam (IB)-induced images of plunge-frozen neurons. The cell body and a few prominent processes can be observed (outlined by white dotted lines). Boxes indicate the regions shown in C and F, respectively. **C, F**: Overlay of fluorescence signal recorded prior to milling with the ion beam-induced image, indicating synapse-rich regions near the cell body that were targeted for cryo-FIB milling. **G-H**: Fluorescence acquired at different stages of cryo-FIB milling (600 nm, 300 nm and 150 nm thickness, respectively). Images show overlays of the fluorescence and reflection channels. While presynaptic Syt1 signal was observable all the way through thin lamellae (G), postsynaptic GluA and GluN signal was generally absent in fine-lamellae (H).

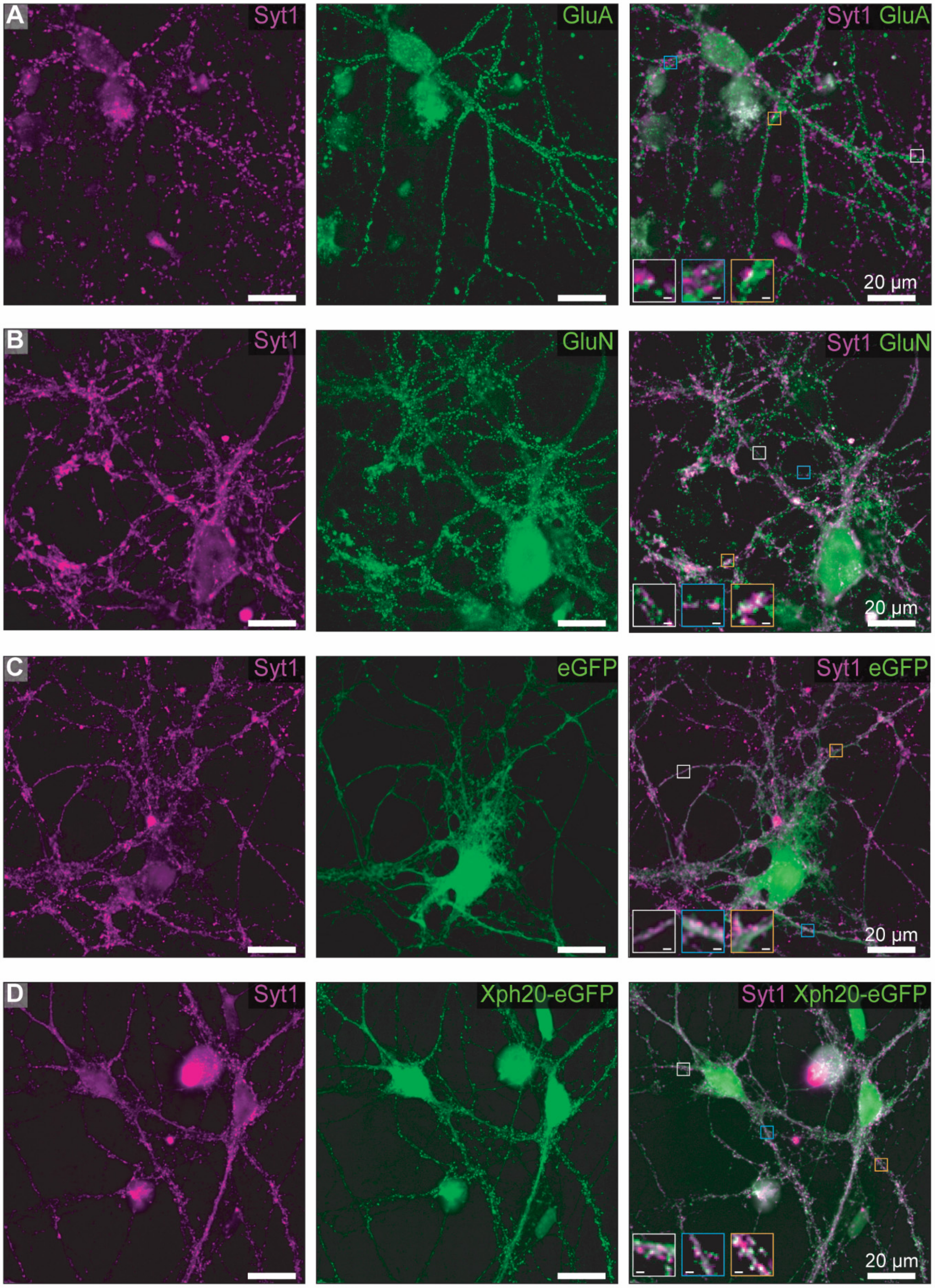


**Figure S3. Room temperature immunofluorescence imaging. A, B**: Imaging of neurons fixed upon live labeling with antibodies marking presynapses (Syt1-ATTO647N, magenta) and postsynapses (A: GluA-ATTO488, green; B: GluN-ATTO565, green). Insets in the merged images highlight co-localized puncta. **C, D**: Imaging of neurons fixed upon live Syt1 labeling (mouse anti-Syt1, anti-mouse-AF647) and AAV transduction with eGFP (C) or Xph20-eGFP (D). In contrast to the diffuse signal of eGFP, Xph20-eGFP showed punctuated signals co-localizing with Syt1 (insets). Scale bars of insets: 1 μm.

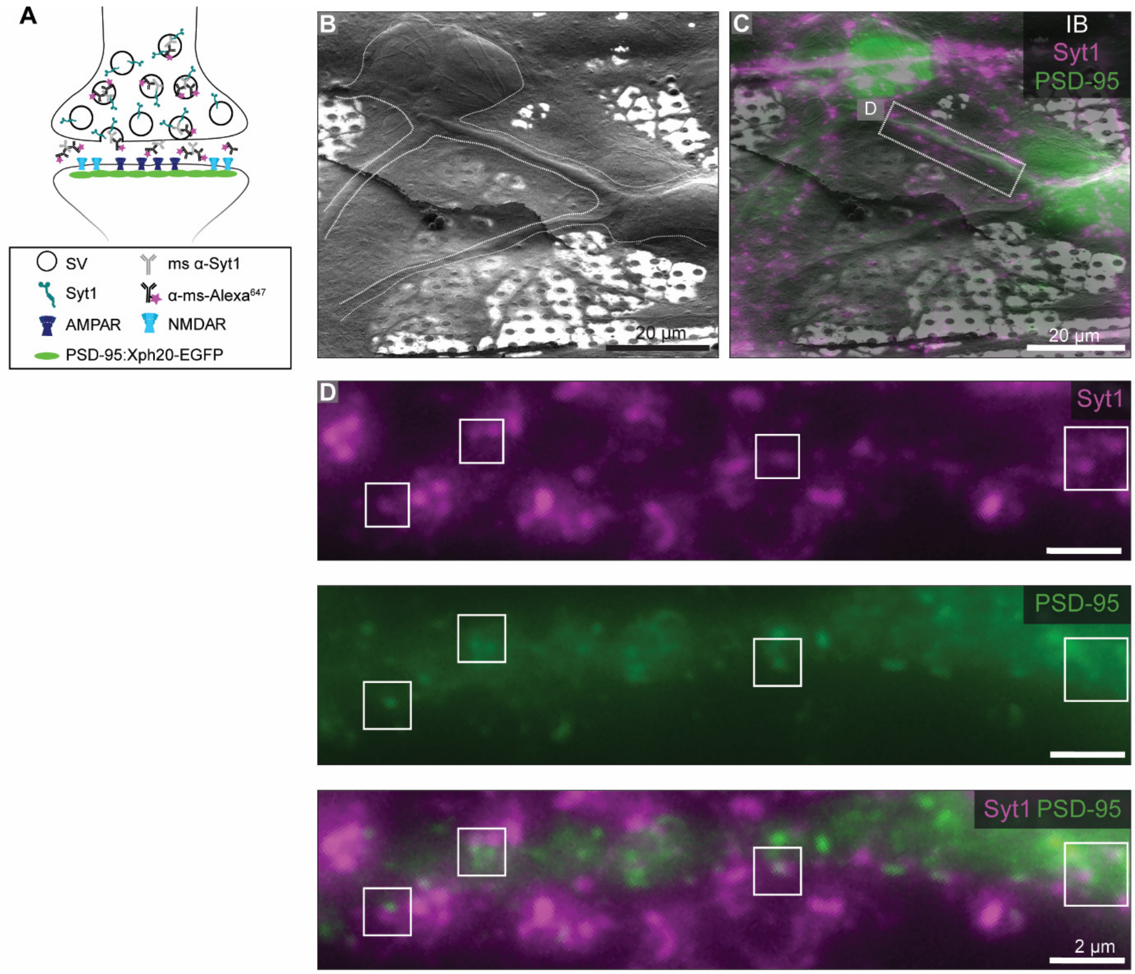


**Figure S4. Combined presynaptic Syt1 immunolabelling and postsynaptic AAV-mediated labelling of PSD-95. A**: Schematic displaying Syt1 antibodies (mouse anti-Syt1, anti-mouse-AF647), AAV-Xph20-eGFP, and their targets in the synapse. **B**: Ion beam (IB)-induced image of two neurons plunge-frozen on an EM grid. **C**: Ion beam-induced image overlaid with fluorescence of Syt1 (magenta) and Xph20-eGFP (green). A box indicates the region shown in (D). **D**: Magnified fluorescence images of pre- and postsynaptic signals along the neuronal process indicated in (C). Boxed regions highlight co-localizations of Syt1 and Xph20-eGFP.

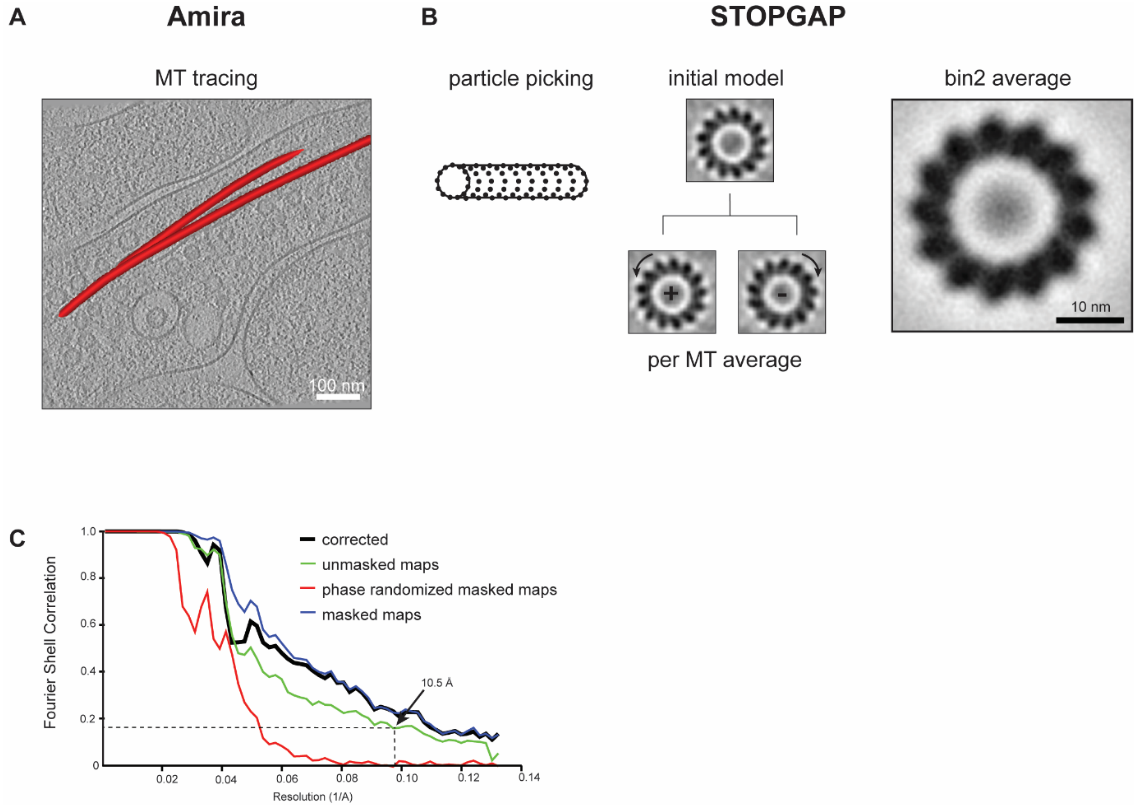


**Figure S5. Workflow for *in situ* subtomogram averaging (STA) of presynaptic microtubules (MTs)**. **A**: Slice through a tomogram showing the tracing of presynaptic MTs represented as tubes (in red) using Amira (Thermo Fisher Scientific). **B**: MT STA workflow in stopgap (Wan et al., 2024). Coordinates obtained from Amira were resampled and plotted on a tube, where each point is placed on a tubulin subunit (left). The initial reference was obtained by alignment and averaging of subtomograms extracted from one MT. All MTs were then aligned against this reference. Individual MT averages (middle) show the polarity of the filament. The polarity of "+" MTs was flipped and subtomograms from all MTs were aligned at bin 4 (7.5 6Å/pixel) and bin2 (3.78 Å/pixel). The right image shows the final bin2 average. **C**: Fourier shell correlation plot of the final average.
